# Dynamics of fixation probability in a population with fluctuating size

**DOI:** 10.1101/2025.08.26.672280

**Authors:** Kavita Jain, Hitesh Sumuni

## Abstract

In many biological processes, the size of a population changes stochastically with time, and recent work in the context of cancer and bacterial growth have focused on the situation when the mean population size grows exponentially. Here, motivated by the evolutionary process of genetic hitchhiking in a selectively neutral population, we consider a model in which the mean size of the population increases linearly. We are interested in understanding how the fluctuations in the population size impact the first passage statistics, and study the fixation probability that a mutant reaches frequency one by a given time in a population whose size follows a conditional Wright-Fisher process. We find that at sufficiently short and long times, the fixation probability can be approximated by a model that ignores temporal correlations between the inverse of the population size, but at intermediate times, it is significantly smaller than that obtained by neglecting the correlations. Our analytical and numerical study of the correlation functions show that the conditional Wright-Fisher process of interest is neither a stationary nor a Gaussian process; we also find that the variance of the inverse population size initially increases linearly with time *t* and then decreases as *t*^−2^ at intermediate times followed by an exponential decay at longer times. Our work emphasizes the importance of temporal correlations in populations with fluctuating size that are often ignored in population-genetic studies of biological evolution.

## I. INTRODUCTION

The remarkable genetic and phenotypic diversity on earth is a product of biological evolution [1] which is a complex process involving several fundamental processes, *viz*., mutation, selection and migration, and various sources of noise [2]. Often these processes act simultaneously and their magnitude and/or direction varies with time; for example, the selection pressure or size of a population may change due to seasonal cycles. While much theoretical work has been done assuming that the variation in selection [3–10] or population size [11–16] is deterministic, in biologically realistic situations, these parameters are random variables. Several recent work have focused on addressing this aspect and assume that the population parameters follow a stochastic process such as a Gaussian process [17–19]; two-state process with the distribution of the switching times between the states to be an exponential (random telegraph process) [20–24] or a power law [25]; and density-dependent birth-death process [26–31].

In the last decade, motivated by the rapid growth of cancer or bacterial cells, the statistical properties of the mutant subpopulation in a population whose size fluctuates in time such that its mean size grows exponentially have been investigated [32–39]. In these studies, the growing population size is modeled as a birth-death process and the mean number of mutants at a genomic site has been analyzed in the framework of a branching process. Here, we are interested in the evolutionary dynamics in a population whose mean size grows *linearly* with time due to genetic hitchhiking in a selectively neutral population [40, 41], as described below.

We consider a finite population of binary sequences with two linked sites where the first site is, in general, under selection and the second one is neutral (that is, the wildtype and mutant type/allele are equally fit). During the evolutionary process, as shown in Fig. 1a, if the mutant allele *A* at the first site escapes loss due to stochastic fluctuations and fixes in the population (that is, it reaches a frequency one), then the frequency of neutral allele 1 initially attached to it also rises and allele 1 thus hitchhikes to fixation [42]. This phenomenon has been shown to result in a decrease in the linked neutral genetic diversity in general models as well as sequence data [43], and was proposed as a mechanism to explain the Lewontin’s paradox [44, 45] which refers to the observation that the neutral genetic diversity seen in natural populations is much smaller than that predicted by the neutral theory [46]. As

**FIG. 1.**
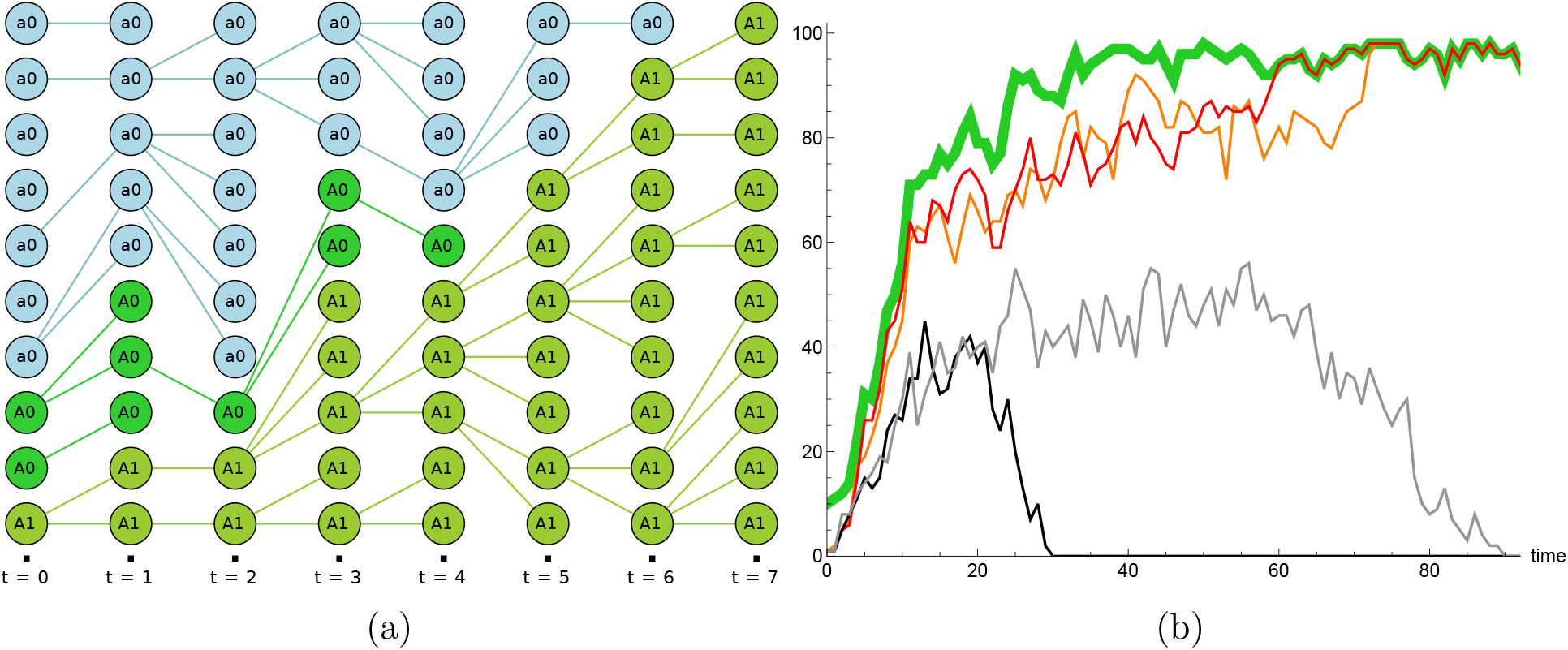
(a) Schematic description of the model depicting Wright-Fisher dynamics in a population of total constant size *N* = 10. At *t* = 0, *N*_0_ = 3 individuals carry *A* allele at the first site and at *t* = 7, *A* fixes in the population. Starting with a single allele 1 (*n*_1_ = 1) in the subpopulation of *A*s, the number of 1s are tracked until the entire population of *A*s is either *A*0 or *A*1 type; note that the allele 1 fixes in the *A* subpopulation at *t* = 5. The effect of the dynamics of *A* allele on the dynamics at the second site can be incorporated as fluctuating population size as described in the text. (b) Sample trajectories of the *A*1 subpopulation (thin lines) for a given trajectory of the *A* allele that eventually fixes (thick line) for *n*_1_ = 1, *N*_0_ = 10, *N* = 100.

Fig. 1a suggests, the dynamics at the second site in the subpopulation of *A*s can be modeled by that of a neutral population of 0s and 1s whose total size is changing with time. If allele *A* is strongly beneficial, the variation in population size can be captured by a deterministic model in which the population size grows exponentially [16] (in related contexts, Gaussian fluctuations about the deterministically growing population have also been considered [47–49]), otherwise the total population size fluctuates and, in our model of interest, its mean size increases linearly at short times.

In this article, we focus on understanding how the time-dependent fixation probability of a mutant is affected due to stochastic changes in the population size. In Sec. II, we define the model and describe the relevant Fokker-Planck equations. The known explxicit solution of these equations when the population size is constant is discussed in Sec. III and a formal solution of the model when the population size fluctuates is given in Sec. IV. Due to temporal correlations in the population size, the model of interest does not seem to be exactly solvable, and therefore we first study the fixation probability ignoring these correlations in Sec. V. Then in Sec. VI, we study a two-point correlation function analytically and higher cumulants numerically which show that the process that describes the changing population size is not a stationary Gaussian process, and discuss how correlations affect the fixation probability. We close the article with a discussion of related models and future directions in Sec. VII.

## II. MODEL

We consider a finite population of binary sequences of length two where the allelic state at the first (second) site can be either *a* or *A* (0 or 1), no mutations between the two alleles at either site occur during the dynamics and all the four sequence configurations are equally fit (but, see Appendix A for a general model). At the first site, the dynamics follow a discrete time, neutral Wright-Fisher process for a population of *constant* size *N* ; in this process, irrespective of the state at the second site, the number 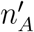 of mutant allele *A* in the current generation is binomially distributed with mean equal to the number *n*_*A*_ of *A*s in the previous generation. Thus the transition probability of the Markov chain that describes this process is given by

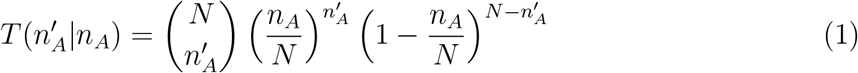

We are interested in the *conditional* process in which, starting from 2 ≤ *N*_0_ *< N*, allele *A* eventually fixes (henceforth referred to as the *A*^*^ process). In this subpopulation of *A*s, we track the number of 1s at the second site whose dynamics also follow the neutral Wright-Fisher process but with changing population size due to the dynamics of allele *A*. Thus, in the current generation, the number of 1s in a subpopulation of size 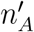 are binomially distributed with mean given by 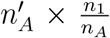 where *n*_1_ is the number of 1s in the previous generation. A schematic illustration of the model and examples of stochastic trajectories of allele 1 while the *A* allele is proceeding to fixation are shown in Fig. 1. Note that the fixation of allele 1 can occur when the population size in the *A*^*^ process is smaller than *N*. We simulated the model described above and obtained the quantities of interest by averaging over 10^3^ −10^4^ fixations of the allele 1 for each trajectory of the *A* allele in 10^3^ −10^4^ independent runs of the *A*^*^ process.

In continuous time, the distribution *P* (*p, t*; *p*_0_, 0) of the frequency 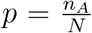 of *A*s at time *t*, starting with frequency 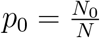 can be described by the following forward Fokker-Planck equation (FPE) [50, 51]:

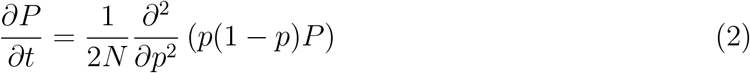

This can be seen by noting that due to Eq. (1), the mean and variance of the change in the frequency *δp* in one generation are, respectively, zero and 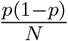, and that the higher moments in *δp* vanish in the scaling limits 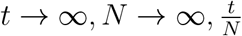 finite. Then from Bayes’ theorem, the above unconditional distribution *P* (*p, t*; *p*_0_, 0) can be related to the distribution *P* ^*^(*p, t*; *p*_0_, 0) conditioned on the fixation of *A* as

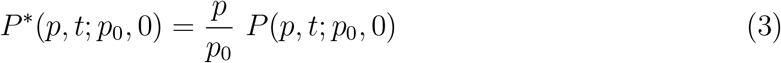

on using that the eventual fixation probability of *A* is equal to its initial frequency *p*_0_ [50, 51] (also see Eq. 9 below). Finally, for a given stochastic trajectory of *A*^*^ process with frequency *p*(*t*), as for Eq. (2), the distribution *X*(*x, t*; *x*_0_, 0) of the frequency 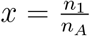 of 1s with initial frequency *x*_0_ obeys the following FPE [16, 51, 52],

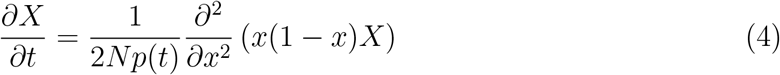

## III. DISTRIBUTIONS FOR CONSTANT POPULATION SIZE

We first describe the known results for the model with constant population size that are pertinent to the discussion here. At *t* → ∞, the subpopulation comprising of *A*s at the first site either goes extinct or gets fixed, but at finite times, the population can either exist in one of these two absorbing states or have a finite frequency 0 *< p <* 1. The solution of Eq. (2) can then be written as [53]

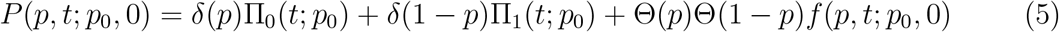

Here, Π_0_(*t*; *p*_0_) and Π_1_(*t*; *p*_0_) are, respectively, the extinction and fixation probability of *A* by time *t*, while *f* (*p, t*; *p*_0_, 0) is the distribution of frequency 0 *< p <* 1 at time *t*.

Due to the conservation of probability for 0 ≤ *p* ≤ 1, it follows from Eq. (2) that the probability current, 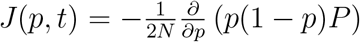 vanishes at *p* = 0 and 1; furthermore, the distribution *P* is normalizable if it diverges weakly enough close to the boundaries. Thus we have [53]

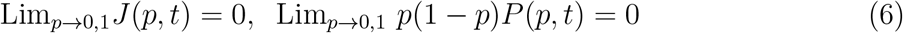

for all *t*. Subject to these boundary conditions, Eq. (2) can be solved using the eigenfunction expansion method on noting that the Jacobi polynomials 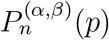 obey (see (18.8.1) of [54]),

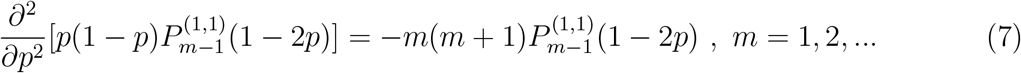

For the initial condition, *P* (*p*, 0; *p*_0_, 0) = *δ*(*p* − *p*_0_), one then obtains [53, 55]

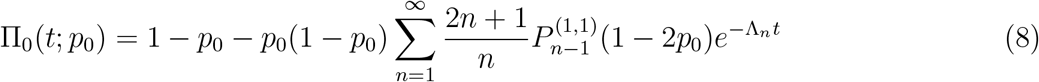

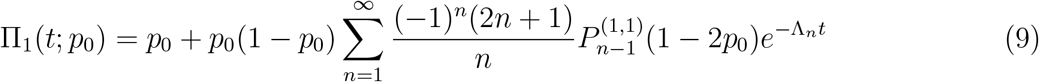

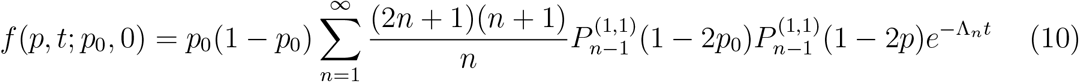

where the eigenvalue,

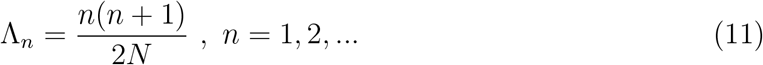

From Eq. (8) and Eq. (9), we note that the probability of eventual extinction and fixation are, respectively, given by 1 − *p*_0_ and *p*_0_.

For numerical purposes, it is convenient to find the absorption probabilities using a backward Fokker-Planck equation given by [50, 51],

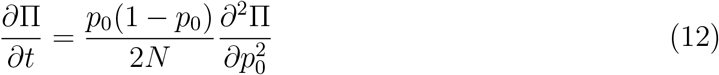

with boundary conditions, Π_0_(*t*; 0) = 1, Π_0_(*t*; 1) = 0 and Π_1_(*t*; 0) = 0, Π_1_(*t*; 1) = 1, and initial condition Π_0_(0; *p*_0_) = Π_1_(0; *p*_0_) = 0 for 0 *< p*_0_ *<* 1.

The simulation results for the dynamics of fixation and extinction probability of *N*_0_ number of *A*s in a population of constant size *N* are shown in Fig. 2a and Fig. 2b, respectively. As intuitively expected, more the initial number of *A*s, higher the chance of fixation and lower the chance of extinction. But while the fixation probability is seen to depend weakly on the initial number of mutants, the extinction probability depends on *N*_0_ and remains negligible on a time scale that increases linearly with *N*_0_. This is because (all) the descendants of all the *A* individuals that are initially present must die for the extinction of allele *A* to occur, but *A* gets fixed if the lineage of any one of the initial *A*s survives.

**FIG. 2.**
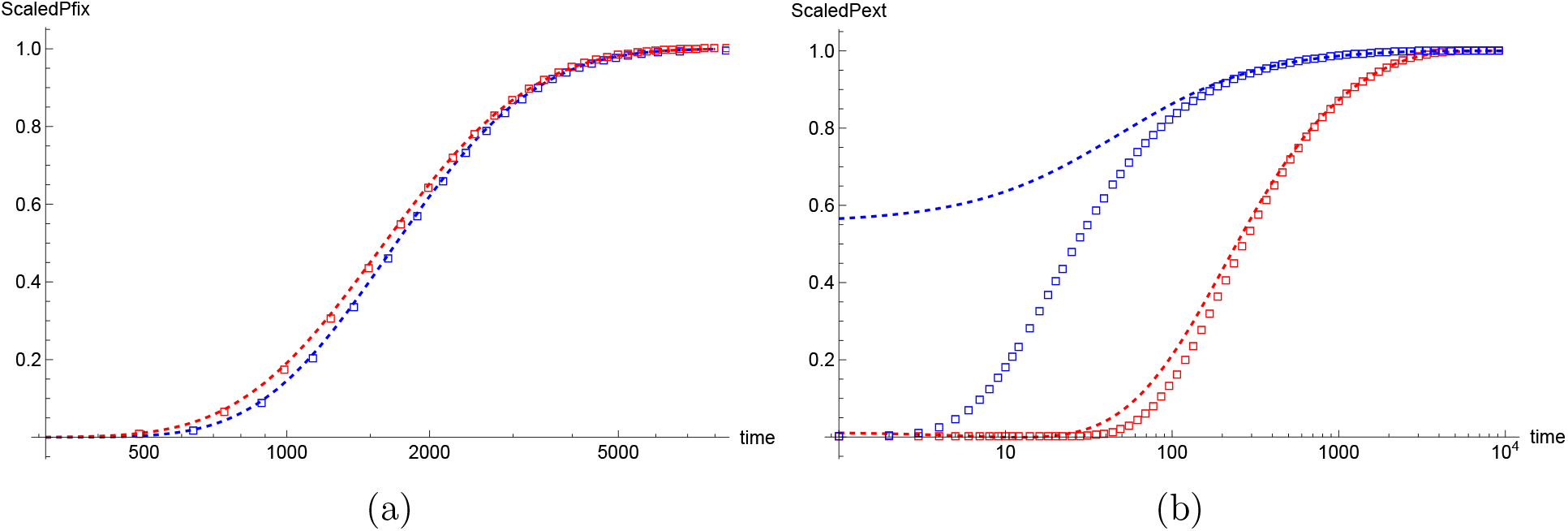
(a) Scaled fixation probability, 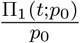 and (b) scaled extinction probability, 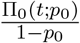 of mutant allele *A* in a population of constant size *N* = 1000, and *Np*_0_ = *N*_0_ = 10 (blue) and 100 (red). The points show the simulation results and the dashed lines are obtained by solving Eq. (12) numerically.

Figure 2 also shows a comparison between the simulation data and the results obtained by numerically solving Eq. (12) with appropriate boundary conditions. We note that while they agree for the fixation probability, there is a strong disagreement for the extinction probability for smaller *N*_0_ at short times - this is because the extinction probability is non-negligible on times of order *N*_0_ but, as explained below Eq. 2, the FPE is valid for large times. An analytical understanding of the relevant time scales predicted by Eq. 8 and Eq. 9 is obtained in Appendix B.

## IV. FIXATION PROBABILITY FOR FLUCTUATING POPULATION SIZE

The probability distribution described by Eq. (4) for changing population size can be obtained in a manner analogous to that for the constant-sized population if one defines the time variable to be 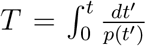. Then, using Eq. (9), the fixation probability, Φ_1_ (*t*; *x*_0_) of the mutant allele 1, starting from frequency *x*_0_, can be written as

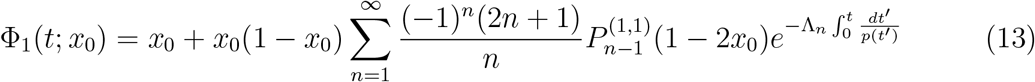

At *t* → ∞, as *p* → 1, it follows from the above equation that even for changing population size, the eventual fixation probability is given by the initial frequency *x*_0_.

On averaging over the fluctuating population size, we obtain ⟨Φ_1_⟩^*^ where the starred angular brackets denote the average with respect to the frequency distribution in the *A*^*^ process. Equation (13) shows that for this purpose, we require [56]

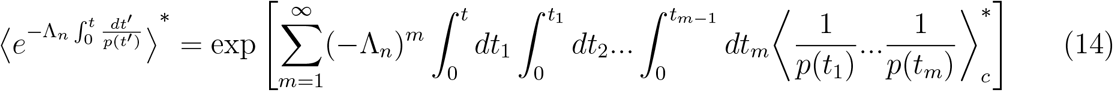

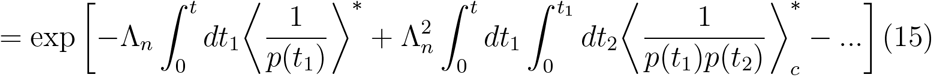

where the subscript *c* denotes the cumulant. If the *A*^*^ process for the inverse frequencies is a stationary Gaussian process, the above expression simplifies considerably (see, for e.g., Eq. (3.75) of [52]). But as shown below, this process is neither stationary nor a Gaussian process, and it does not appear possible to obtain an exact expression for the fixation probability in the full model. Therefore, in Sec. V, we first consider a model that ignores all the temporal correlations, and then study the nature of the correlations and their effect on fixation probability in Sec. VI.

Figure 3a shows the simulation results for the fixation probability of a single mutant 1 when the population size is changing in the full model defined in Sec. II and in the uncorrelated model described in Sec. V, and when the population size remains fixed at the initial size *N*_0_. We note that until time of order *N*_0_, the fixation probability is almost the same in the three models. But at larger times *t* ≫ *N*_0_, the mutant allele 1 is more likely to fix when the population size remains constant at *N*_0_ than when it is increasing as it is harder to fix in a larger population. Furthermore, as displayed in Fig. 3b, at times of order *N*, the fixation probability in the full model can be approximated by that in the uncorrelated model. However, as Fig. 3a shows, at intermediate times, fixation is much more likely in the uncorrelated model than in the full model; see Sec. VI C for a discussion.

**FIG. 3.**
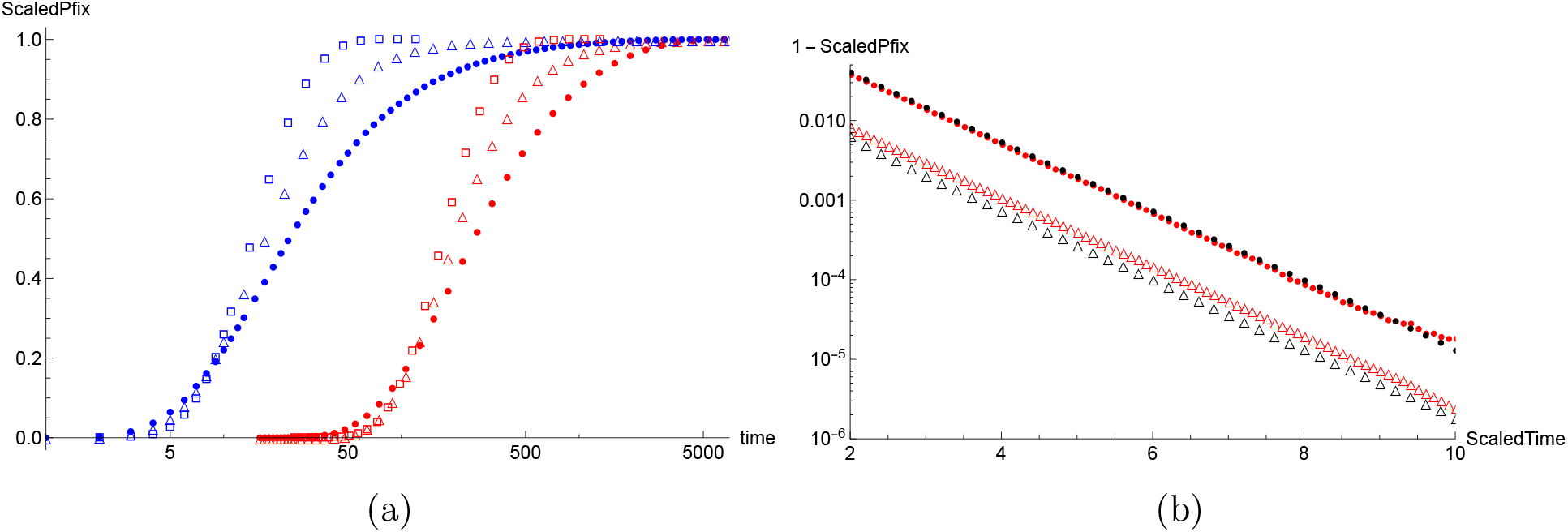
(a) Fixation probability of a single mutant allele 1 in a subpopulation of *A*s with initial size *N*_0_ = 10 (blue) and 100 (red) divided by the initial frequency, 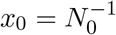. The simulation data for the full model (•) and uncorrelated model (△) for *N* = 1000, and when population size remains constant at the initial size (□) are shown. (b) Figure shows the complementary scaled fixation probability as a function of scaled time, 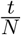 for *N*_0_ = 100, *N* = 1000 (red) and *N*_0_ = 50, *N* = 500 (black) for full model (•) and uncorrelated model (△).

## V. UNCORRELATED POPULATION SIZE

In this section, we assume that the inverse frequencies in the *A*^*^ process are uncorrelated in time. Then Eq. (14) gives

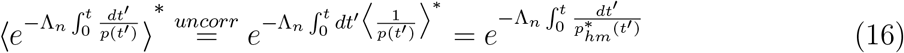

where, 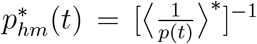 is the time-dependent, conditional harmonic mean frequency, and is given by

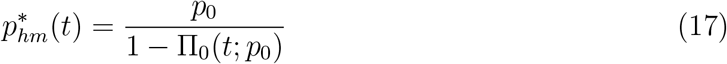

as shown below [see Eq. (20)].

### A. Conditional mean and harmonic mean

We now discuss how the arithmetic mean and harmonic mean of frequency *p* change with time. Evidently, these quantities are identical in the full model and the uncorrelated model. We first consider the (arithmetic) mean frequency in the *A*^*^ process which is given by 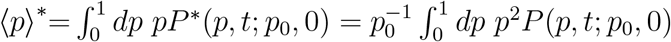 due to Eq. (3). Then from Eq. (2), we obtain

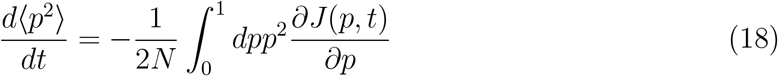

where, 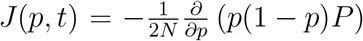 is the probability current. On integrating the RHS of the above equation by parts and using the boundary conditions given in Eq. (6), we obtain an equation for the dynamics of ⟨*p*⟩^*^ which can be easily solved, and we find that

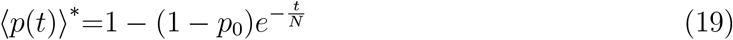

The above expression shows that the mean frequency in the *A*^*^ process grows *linearly* at short times (*t* ≪ *N*); this is unlike when the *A* allele is under positive selection and its conditional mean frequency rises exponentially [16]. Note that the time scale *N*_0_ does not appear in the dynamics of the arithmetic mean frequency.

The harmonic mean frequency conditioned on fixation is given by Eq. (17) since

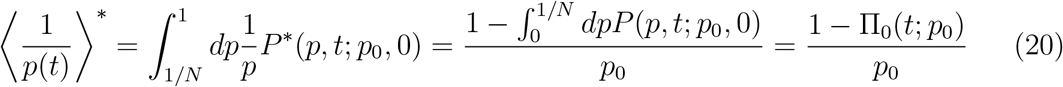

on using that the distribution *P* (*p, t*; *p*_0_, 0) is normalized to one, and due to Eq. (5). Since Π_0_(0; *p*_0_) = 0 and Π_0_(∞; *p*_0_) = 1 − *p*_0_, it follows from Eq. (20) that the harmonic mean frequency eventually reaches one, starting from *p*_0_. To understand the dynamics of the conditional mean of the inverse frequency, we use the results in Appendix B and find that

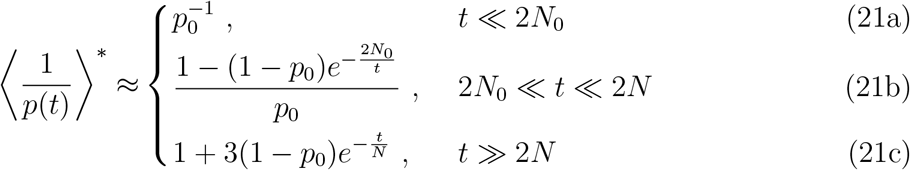

The above expression shows that the conditional harmonic mean frequency remains constant on a time scale proportional to the initial number of mutants in the *A*^*^ process and approaches its asymptotic value on a time scale that grows linearly with the total population size *N*.

Figure 4a shows that Eq. (17) and Eq. (19) agree with the corresponding results obtained using numerical simulations, and as expected, the conditional mean frequency is larger than the corresponding harmonic mean frequency. The harmonic mean frequency obtained using the approximations in Eq. (21) is also compared with the simulation results, and we find a reasonable agreement in their respective regimes of validity.

**FIG. 4.**
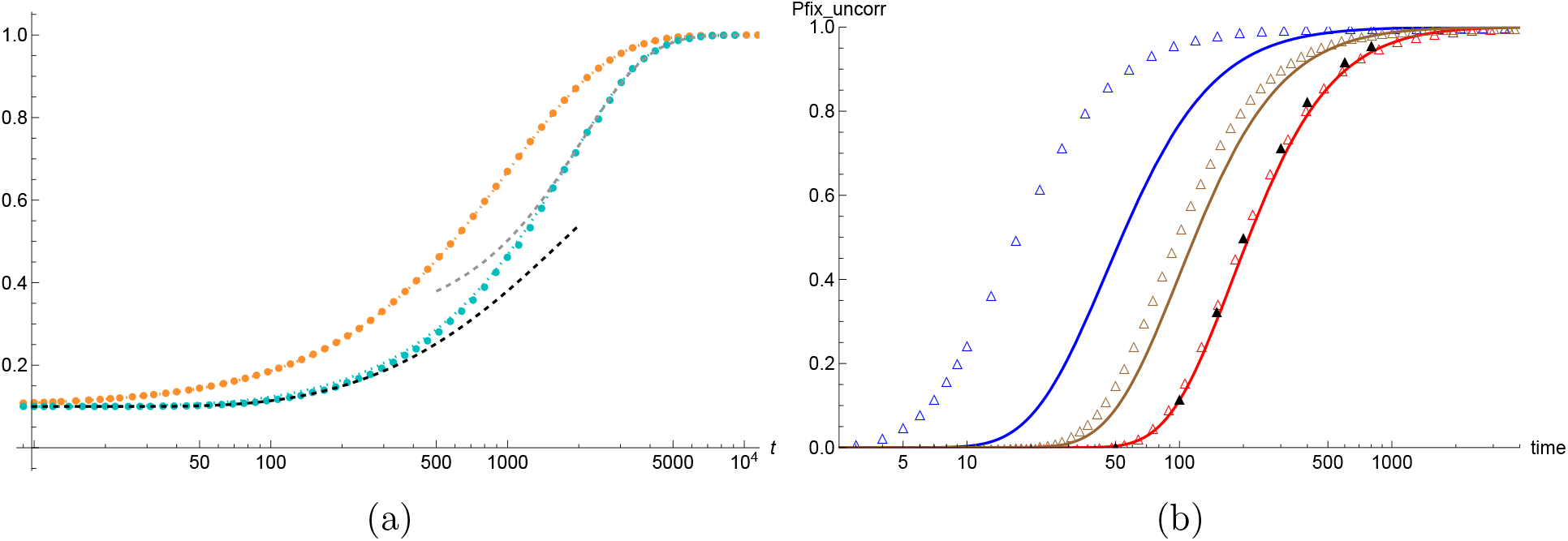
(a) Arithmetic mean frequency (orange) and harmonic mean frequency (cyan) in the *A*^*^ process obtained using numerical simulations (points) are compared with the exact expressions (solid lines) given by Eq. (19) and Eq. (17), respectively, for *N* = 1000, *p*_0_ = 0.1. The RHS of Eq. (17) is obtained by numerically integrating Eq. (12) with appropriate boundary conditions. The approximate expressions for the conditional harmonic mean frequency obtained using Eq. (21b) and Eq. (21c) are shown by black and gray dashed lines, respectively. (b) Scaled fixation probability, 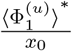 of a single mutant allele 1 in the uncorrelated model for *N* = 1000, and 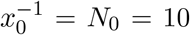 (blue), 50 (brown) and 100 (red) obtained from simulations (△) and numerically solving Eq. (23) (lines) is shown. The approximate result (▲) given by Eq. (24) is shown for *N*_0_ = 100.

### B. Fixation probability

From Eq. (13) and Eq. (16), we obtain the fixation probability averaged over the uncorrelated population size to be

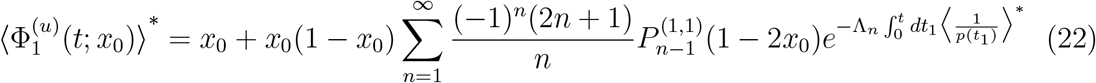

Thus, similar to Eq. (12) for constant population size, the above fixation probability is a solution of the following backward FPE,

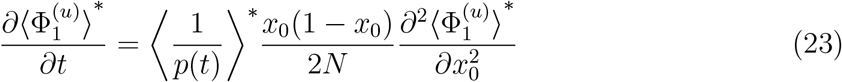

with boundary conditions, 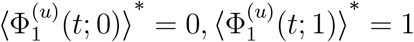, and initial condition 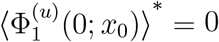; the results obtained by numerically integrating the above equation are shown in Fig. 4b. We also measured the probability 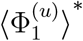 in simulations in which a new stochastic trajectory of the subpopulation *A* that eventually fixes was generated at *each* generation of the Wright-Fisher process for the mutant allele 1 so that the subpopulation size at all times are uncorrelated. As shown in Fig. 4b, the simulation results for small *N*_0_ are not captured by Eq. (23) for reasons already discussed in Sec. III.

But for sufficiently large *N*_0_, on using Eq. (21b) for *t* ≪ 2*N* in Eq. (22), we obtain

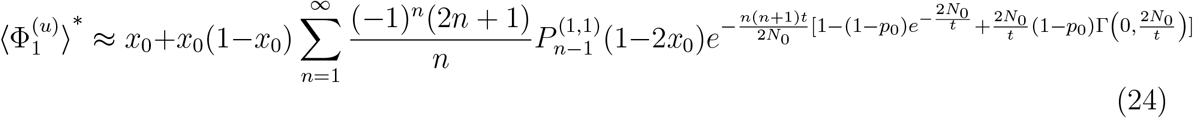

which is in good agreement with the simulation results shown in Fig. 4b. At longer times *t* ≫ 2*N*, the dynamics are determined by the smallest eigenvalue Λ_1_, and as shown in Appendix C, the fixation probability approaches its asymptotic value as

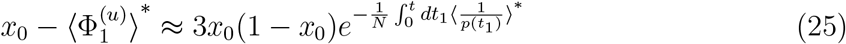

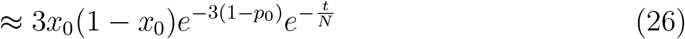

As expected and in agreement with the simulation results in Fig. 3b, the approach to the eventual fixation probability occurs over a time scale *N*. The above equations also show that the amplitude of the exponential decay depends on both *p*_0_ and *x*_0_, and the simulation data shown in Fig. 3b for two values of *x*_0_ is consistent with the above prediction.

## VI. CORRELATIONS IN THE *A*^*^ PROCESS

As Fig. 3a and Fig. 3b show, the uncorrelated population size model does not accurately capture the fixation probability in the full model, and is a particularly poor approximation at intermediate times. Thus the temporal correlations in Eq. (14) can not be ignored; below we study these correlations numerically and analytically using FPEs for sufficiently large *N*_0_, and also discuss how they affect the fixation probability.

### A. Two time correlation function

We first consider the unconnected two-time correlation function for the inverse population frequency in the *A*^*^ process. For *t*_2_ ≥ *t*_1_, we can write

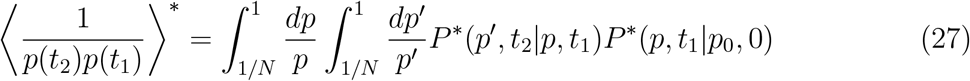

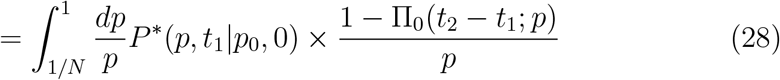

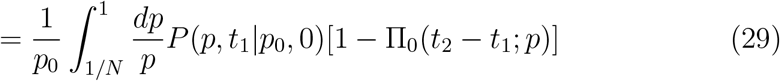

on using Eq. (3) and Eq. (20), and where *P* (*p, t*_1_|*p*_0_, 0) is given by Eq. (5). Due to the nonzero lower limit of integration, the first term on the RHS of Eq. (5) does not contribute to the above integral. Then due to the second term on the RHS of Eq. (5) and the boundary condition Π_0_(*t*; 1) = 0, we obtain

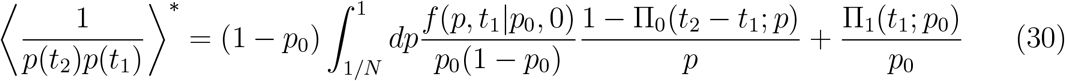

Using the above equation and Eq. (20), the connected correlation function

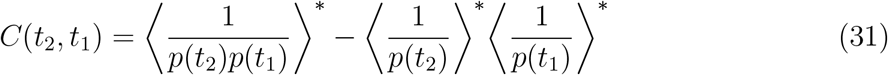

can be obtained.

#### 1. Variance of the inverse frequency

For *t*_1_ = *t*_2_ = *t*, due to the initial condition Π_0_(0; *p*_0_) = 0, Eq. (30) reduces to

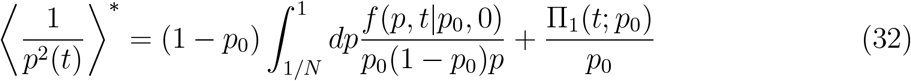

For *t* ≪ 2*N*, as explained in Appendix B, the fixation probability Π_1_ ≈ 0 so that the last term on the RHS of the above equation can be ignored. Furthermore, as shown in Appendix D, for *p*_0_ → 0, *t* ≪ 2*N*,

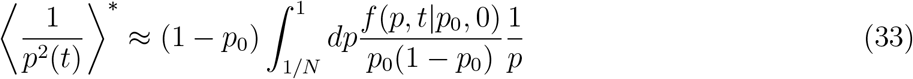

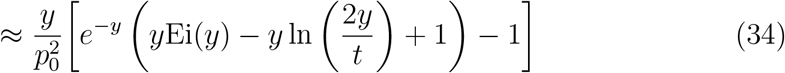

where 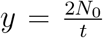 and 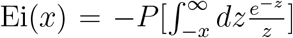 is the exponential integral [see (6.2.6) of [54]]. For *t* ≪ 2*N*_0_, using (6.12.2) of [54] for the asymptotic expansion of Ei(*x*), we find that the RHS of Eq. (34) can be approximated by 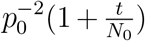. But for 2*N*_0_ ≪ *t* ≪ 2*N*, using (6.6.1) of [54] for the power series expansion of Ei(*x*), we obtain 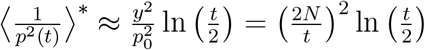. Thus, at intermediate times, the second moment of the inverse frequency in the *A*^*^ process decays algebraically and is independent of *N*_0_. At larger times where *t* ≫ 2*N*, using the results in Appendix B and Appendix D, we obtain 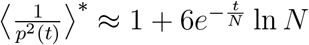.

Using the above approximations for the second moment and Eq. (21) for the mean of the inverse frequency, we find that the dynamical behavior of the variance falls in three distinct regimes: initial linear increase, algebraic decay as *t*^−2^ and finally an exponential approach to zero:

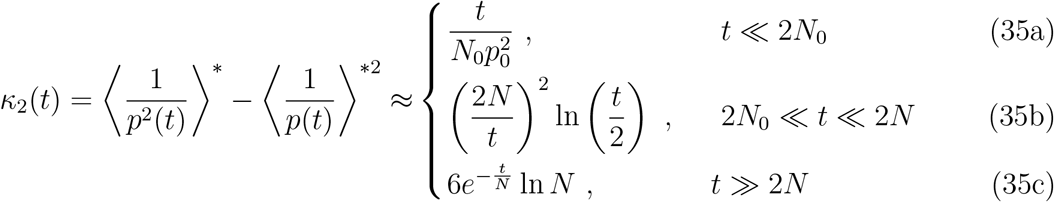

The above expression also shows that the maximum in the variance occurs at time ∼ *N*_0_. Figure 5a shows a comparison between the simulation results and the above approximations for the variance. We find that for *t* ≪ 2*N*, the variance obtained using Eq. (34) and Eq. (21b) is in good agreement with the simulations; the initial linear growth indicated by Eq. (35a) is also observed but the approximate expression given by Eq. (35b) at intermediate times is seen for a very short time span as the exponential decay given by Eq. (35c) sets in.

**FIG. 5.**
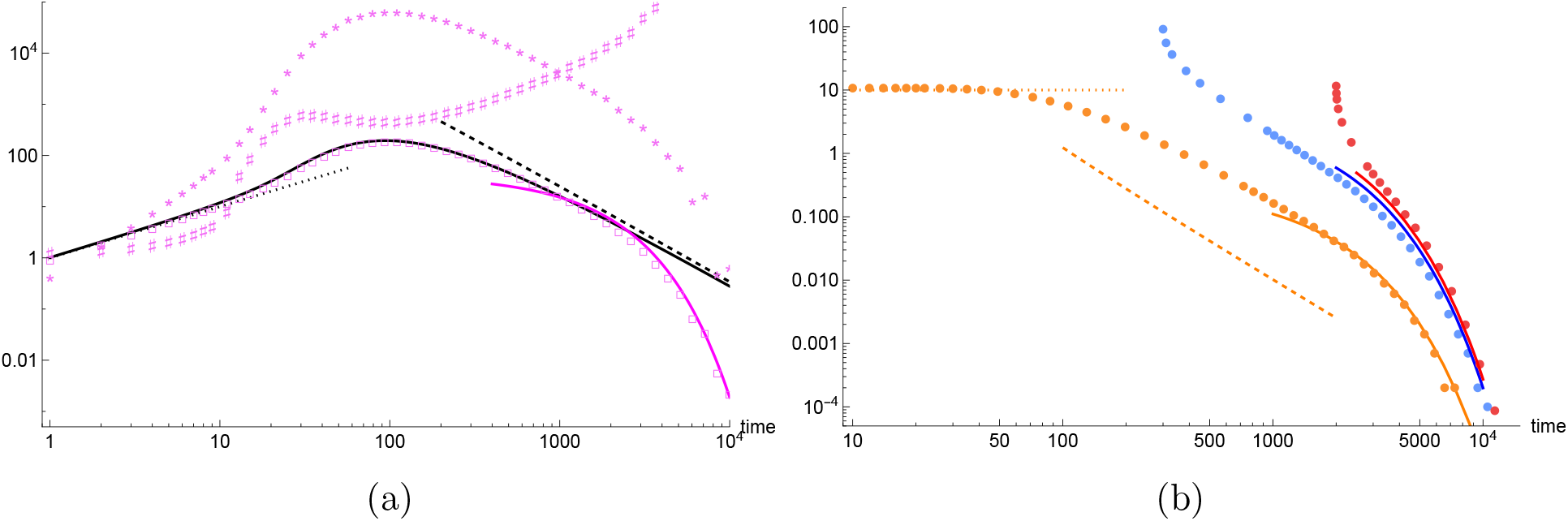
(a) Variance (□), skewness (⋆) and scaled kurtosis (*♯*) of the inverse frequency in the *A*^*^ process as a function of time for 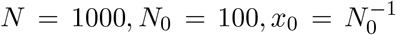 obtained from simulations is shown. The solid lines show the variance obtained using Eq. (34) and Eq. (21b) for *t* ≪ 2*N* and Eq. (35c) for *t* ≫ 2*N* ; the approximate expressions Eq. (35a) and Eq. (35b) are depicted by dotted and dashed line, respectively. (b) Unequal time correlation function, 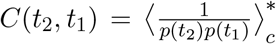 as a function of time *t*_2_ for *t*_1_ = 10 (orange), 300 (blue), 2000 (red) is obtained from simulations (points). The dotted line shows Eq. (35a) evaluated at time *t*_1_ = 10 and the dashed line shows 0.01(*t*_2_ − *t*_1_)^−2^ to test the scaling in Eq. (37) for *t*_1_ = 10; the solid lines show Eq. (38) in their respective regions of validity.

#### 2. Unequal time correlation function

For *t*_2_ *> t*_1_, we first note that, as discussed in Sec. III, Π_0_(*t*_2_ −*t*_1_; *p*) ≈ 0 for *t*_2_ −*t*_1_ ≪ 2*Np*. Then for a given *t*_1_ ≪ 2*N*_0_, the distribution *f* (*p, t*_1_|*p*_0_, 0) is expected to remain close to the initial frequency *p*_0_, and therefore, for *t*_1_ ≪ 2*N*_0_, *t*_2_ − *t*_1_ ≪ 2*N*_0_, the unequal time correlation function given by Eq. (30) can be approximated by the second moment at time *t*_1_ [see Eq. (32)]. But at larger times (*t*_2_ − *t*_1_ ≫ 2*N*_0_) where the extinction probability Π_0_ is non-negligible, the correlation function depends on time *t*_2_ also. The simulation data in Fig. 5b for *t*_1_ = 10 shows that indeed *C*(*t*_2_, *t*_1_) ≈ *κ*_2_(*t*_1_) when *t*_2_ ≪ 2*N*_0_ and decreases thereafter.

Then, as shown in Appendix E, except when both *t*_2_, *t*_1_ ≲ 2*N*_0_, the unconnected two-time correlation function defined in Eq. (30) can be rewritten as

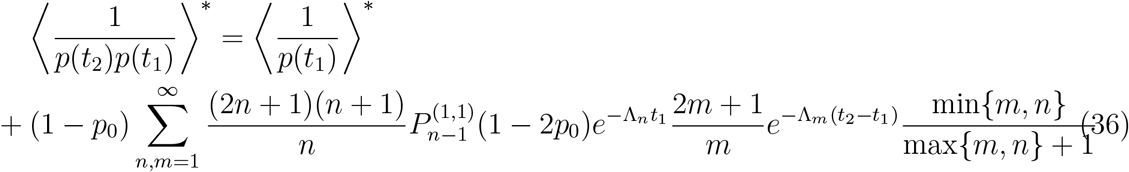

As for variance, due to the exponential factors in the above summand, one can consider cases when *t*_1_ and *t*_2_ − *t*_1_ are smaller or larger than 2*N* to obtain approximate expressions for the unequal time correlation function.

We first consider the parameter regime where *t*_1_ ≪ 2*N*_0_ but 2*N*_0_ ≪ *t*_2_ − *t*_1_ ≪ 2*N*. Then, as discussed in Appendix F, we obtain

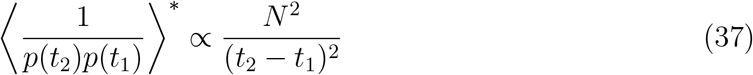

which is in reasonable agreement with the simulation results shown in Fig. 5b for *t*_1_ = 10; a better quantitative agreement seems difficult to obtain due to the crossovers on either side of the intermediate regime, as mentioned above for the variance. For *t*_1_ ≪ 2*N*_0_ and *t*_2_ − *t*_1_ ≫ 2*N*, see Eq. (38a) below.

For *t*_1_ ≫ 2*N*_0_, as Fig. 5b shows, much of the dynamics of the correlation function can be captured by its late time behavior where *t*_2_ − *t*_1_ ≫ 2*N*. Then as described in Appendix G, we obtain

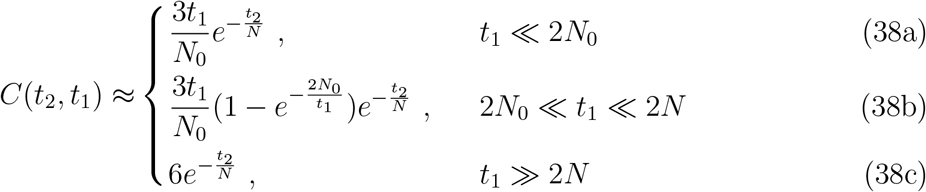

These expressions are found to be in good agreement with the simulation results shown in Fig. 5b for various *t*_1_.

### B. Skewness and kurtosis

As described in the last subsection, the correlation function depends not only on the time difference but also on the earlier time *t*_1_, and therefore the *A*^*^ process for the inverse frequency is not a stationary process. We now ask if this process is a Gaussian process; here, we do not address this question analytically and instead numerically measure the skewness and scaled kurtosis defined as

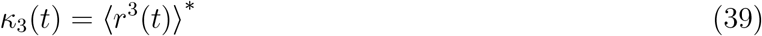

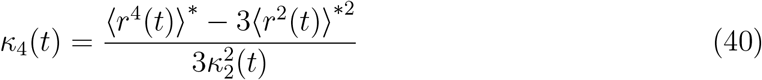

where, 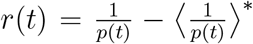. For a stationary Gaussian process, *κ*_3_(*t*) = 0, *κ*_4_(*t*) = 1. But as the simulation results in Fig. 5a show, *κ*_3_(*t*) is nonzero, and, as for variance, it is also a nonmonotonic function of time with an algebraic decay at intermediate times. The scaled kurtosis is seen to be close to one at short times but it is also a nonmonotonic function of time. As the kurtosis must vanish at large times when the allele *A* has fixed, an increase in *κ*_4_ for *t* ≳ 1000 is likely a numerical artefact. We thus conclude that the *A*^*^ process for the inverse frequency is not a Gaussian process.

### C. Effect of correlations on the fixation probability

We first note that as displayed in Fig. 3a for *N*_0_ = 100, the fixation probability in the full model differs from that in the uncorrelated model for 100 ≲ *t* ≲ 1000 which, as Fig. 5 shows, is also the time range where variance and unequal time correlation function *C*(*t*_2_, *t*_1_) for *t*_1_, *t*_2_ ≪ 1000 decay algebraically. Thus, in general, we expect that the fixation probability in the full model can not be approximated by that in the uncorrelated model at intermediate times where 2*N*_0_ ≪ *t* ≪ 2*N*.

Furthermore, Fig. 3a also shows that in the full model, the fixation probability on these time scales is *smaller* than that in the uncorrelated model suggesting that the population size is effectively larger in the former case. To understand this, we measured the probability *Q*(*N*_*f*_) that the population size in the *A*^*^ process is *N*_*f*_ at the time when allele 1 fixes. As Fig. 6a shows, this distribution is heavily skewed towards small *N*_*f*_ in the uncorrelated model; however, this does not mean that when correlations are neglected, the fixation occurs at very small times as the distribution of fixation time in the uncorrelated model and full model are quite similar, see the inset of Fig. 6a. Rather when correlations are absent, the population size fluctuates substantially between generations and therefore, even at large times, its size can be small (although the mean size increases according to Eq. (19)). In contrast, due to correlations in the full model, the population size increases in a smoother fashion and allele 1 typically encounters a large population (see Fig. 6a).

**FIG. 6.**
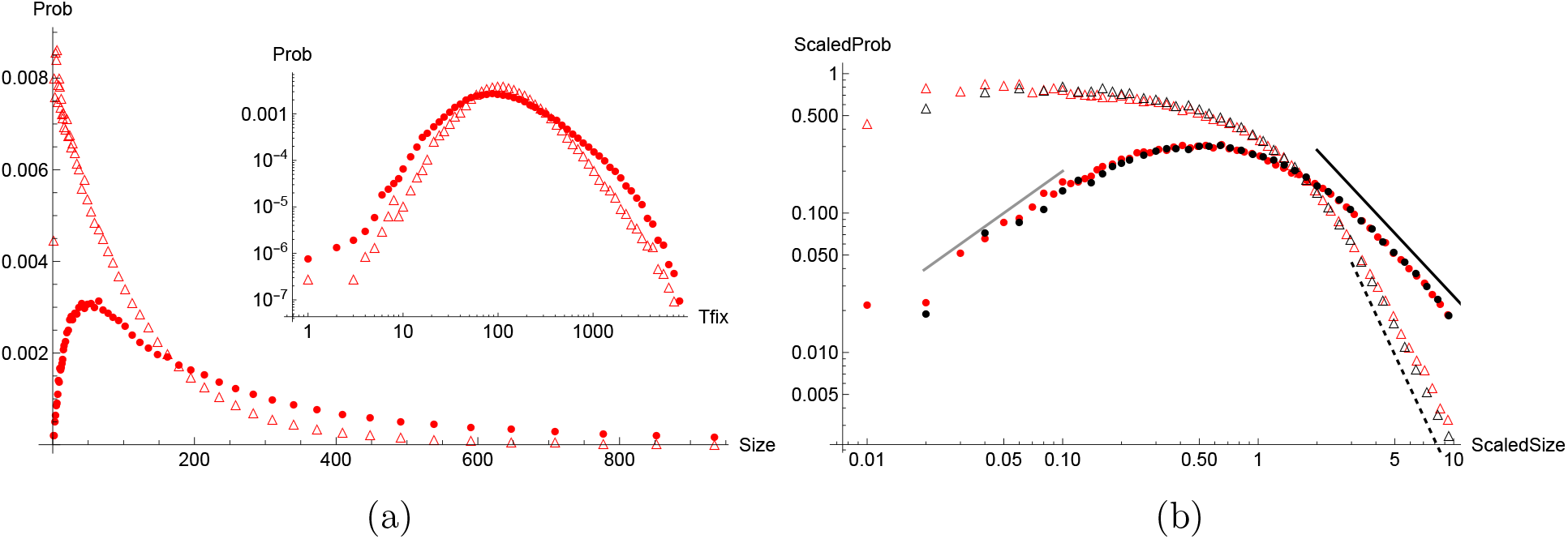
(a) Distribution of population size *N*_*f*_ at the time of fixation of allele 1 (main) and distribution of fixation time (inset) for *N* = 1000, *N*_0_ = 100 obtained from simulation for the full model (•) and uncorrelated model (△). (b) Scaled distribution, *N*_0_*Q*(*N*_*f*_) as a function of scaled size, 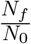 for *N* = 1000, *N*_0_ = 100 (red) and *N* = 500, *N*_0_ = 50 (black) for the full model (•) and uncorrelated model (△). The slope of the solid lines are 1 (gray) and −3*/*2 (black), and that of dashed line is −3.

Figure 6b suggests that for large *N*_*f*_, the distribution *Q*(*N*_*f*_) decreases algebraically, as 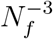 in the uncorrelated model and 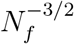 in the full model. The behavior in the former case can be obtained analytically as explained in Appendix H, and we find that the population size distribution is of the following scaling form,

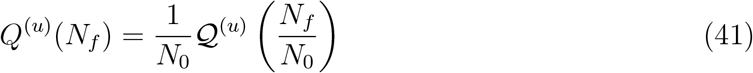

where, the scaling function, 𝒬 ^(*u*)^(*x*) is a constant for *x* ≪ 1 and decays as *x*^−3^, *x* ≫ 1. As the data collapse in Fig. 6b indicates, the above scaling form holds for the full model also but the scaling function changes nonmonotonically; a numerical fit suggests that the scaling function initially increases as *x* and then decays as *x*^−3*/*2^.

## VII. DISCUSSION

Natural populations have a finite carrying capacity, but their size does not remain fixed at the maximum possible size and, in general, it changes stochastically in a correlated fashion. However, most work in population-genetic studies of biological evolution implicitly assume that the population sizes are uncorrelated, and subsume the effect of changing size in an effective population size [57] given by the harmonic mean of the varying population size (see Eq. (17)); however, an effective population size does not exist when the correlations can not be neglected [20, 24, 58]. In a similar vein, here we have shown that the chance of fixation is strongly affected during the time regime when correlations in the population size are substantial.

Unfortunately, we are unable to obtain exact expression for the time-dependent fixation probability as the *A*^*^ process which is the conditional Wright-Fisher process of our interest is found to be neither a stationary nor a Gaussian process; an example where exact results for the dynamics of fixation probability have been obtained is given in [24] where the population size follows a random telegraph process, which is a stationary process and where the correlations in the inverse population size decay exponentially. However, such analytically tractable models make adhoc assumptions for the variation in the population size [12, 14, 23, 24, 29], while the stochastic process governing the fluctuations in the population size considered here arises naturally from a well established model of genetic hitchhiking [42].

Here, we have discussed the dynamics of fixation probability when the *A* subpopulation is neutral. But when the subpopulation is under selection (and therefore, grows exponentially), although the eventual fixation probability has been studied [27, 30, 31], to our knowledge, the dynamics have not been investigated and should be addressed. In this work, the analytical results are obtained when the initial size of the *A* subpopulation is of the order of the total population size because the Fokker-Planck equations analyzed here do not hold otherwise (see Fig. 2b and Fig. 4b). However, in biologically relevant situations, the initial size *N*_0_ ∼ 𝒪 (1) and one needs to work with the discrete number of individuals (and not continuous frequency) at short times; a detailed analyses, perhaps along the lines of [29], is desirable.

## Acknowledgments

HS thanks JNCASR for support through the SRFP-2024 and DST for funding through the INSPIRE-SHE, and is grateful to Venu Goswami, Ayantik Kundu, Prashant Singh and Lakshita Jindal for helpful discussions.

## Appendix A: General model

To understand how other sites linked to selectively neutral regions of the genome affect the neutral genetic diversity [43], one considers a finite population of binary sequences denoted by {*σ*_1_, …, *σ*_*L*_; *η*_1_, …, *η*_*ℓ*_} where *σ*_*i*_ = *a, A* and *η*_*i*_ = 0, 1, and *L, ℓ* ≫ 1. It is usually assumed that (i) the sites labeled by *σ*_*i*_ are, in general, under selection and the *η*_*i*_-sites are neutral, (ii) for long sequences and low mutation rates, at most one mutation occurs at a site from wildtype allele (*a* or 0) to mutant allele (*A* or 1) and the reverse mutations can be neglected [59], and (iii) the linkage between the sites can be broken due to genetic recombination; for a recent work on models that incorporate such and other biological details, see [16] and references therein. Here, as we are mainly interested in understanding the effect of fluctuating population size on the fixation probability of a mutant at a fully linked neutral site, we have focused on a model with *L* = *ℓ* = 1 and assumed that all the four sequence configurations are equally fit.

## Appendix B: Absorption probabilities in constant-sized population

Here, we analyze the extinction and fixation probability in a population of fixed size *N*. First, consider the sum on the RHS of Eq. (8) for extinction probability,

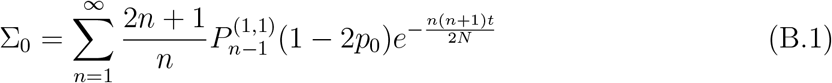

on using Eq. (11). For *t* ≫ 2*N*, it is sufficient to consider the term corresponding to the lowest eigenvalue (*n* = 1) which yields

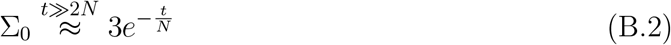

since the Jacobi polynomial 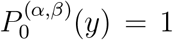. On the other hand, for *t* ≪ 2*N, n* ≫ 1 also contribute to the sum Σ_0_. Then it is useful to approximate the Jacobi polynomials as [16]

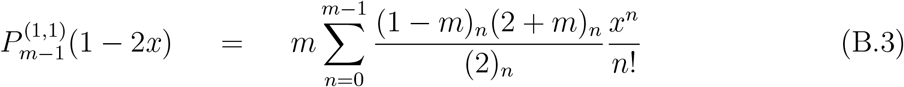

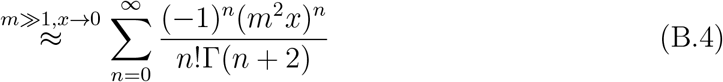

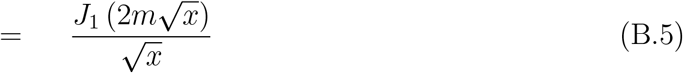

which is obtained in the scaling limit 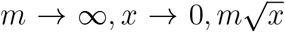 finite, and where *J*_1_(*y*) is the Bessel function of the first kind. For small *p*_0_, we can then write

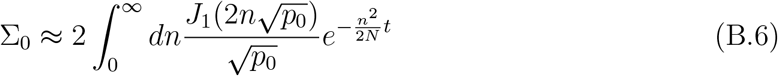

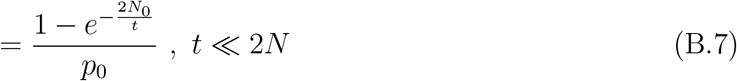

which is valid when *t, N*_0_ ≫ 1 and 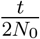 are finite. Thus, as discussed in Sec. III, the results obtained from the FPE agree with those from simulations if the initial number of mutants and time are sufficiently large. Equation (B.7) also shows that for *t* ≪ 2*N*_0_, as 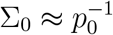, the extinction probability Π_0_ in Eq. (8) is negligible; furthermore, using Eq. (B.7) in Eq. (20), we obtain Eq. (21a) and Eq. (21b), and similarly, Eq. (B.2) leads to Eq. (21c).

Next, consider the sum on the RHS of Eq. (9) for fixation probability,

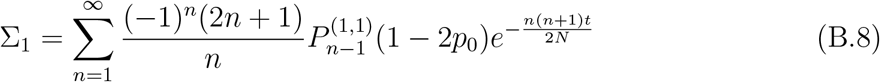

As for Σ_0_, for *t* ≫ 2*N*, it is sufficient to keep the term corresponding to the lowest eigenvalue in the above sum, and we have 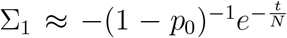. But even at shorter times where *t* ≪ 2*N*, our numerical study of the above sum shows that the alternating series Σ_1_ converges rapidly and can be approximated by first few terms in the summand of Σ_1_. We also find that the sum remains close to its initial value −(1 − *p*_0_)^−1^ until *t* ≲ *N*, in accordance with the conclusion in the main text that the initial number *N*_0_ does not play a role in the dynamics of Π_1_.

## Appendix C: Long time dynamics in the uncorrelated model

For the uncorrelated model, on retaining only the term corresponding to the lowest eigen-value in Eq. (22), we get

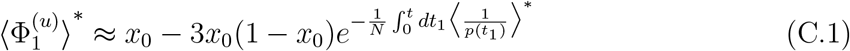

since 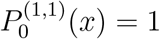. Then, due to Eq. (8), we obtain

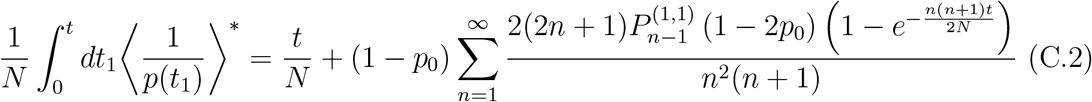

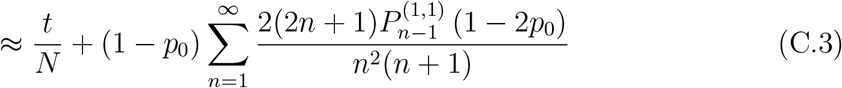

so that

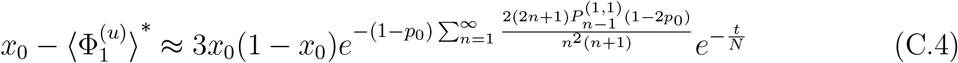

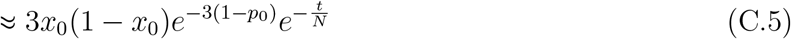

## Appendix D: Approximations for the variance of the inverse frequency

Consider the integral on the RHS of Eq. (32) which can be written as

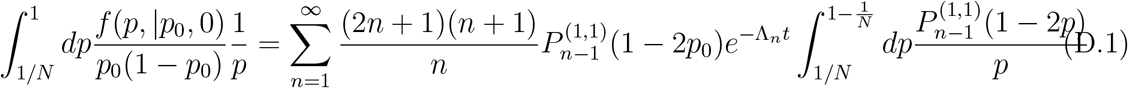

on using Eq. (10). We first note that for *t* ≪ 2*N*, most of the contribution to the integral on the RHS comes from *p* → 0 and to the sum from *n* ≫ 1. Then using the approximation in Eq. (B.5) for the Jacobi polynomial in the above equation and on approximating the sum by an integral, we obtain

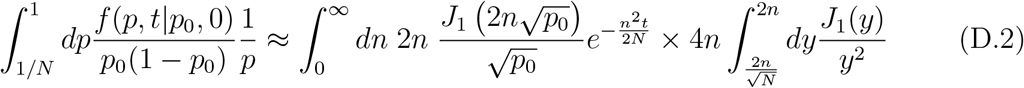

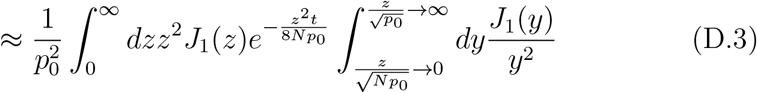

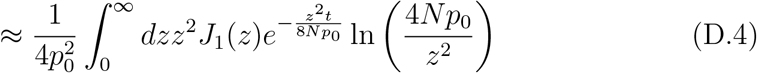

where the last expression is obtained on using that the lower limit in the inner integal in Eq. (D.3) scales as 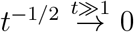 while the upper limit 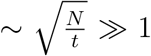 as we are working in the regime where *t* ≪ 2*N*. On performing the integral in Eq. (D.4), we finally obtain Eq. (34) in the main text. For *t* ≫ 2*N*, it is sufficient to evaluate the sum on the RHS of (D.1) with *n* = 1 which yields 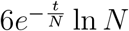.

## Appendix E: Unequal time correlation function

Consider the integral on the RHS of Eq. (30) when either *t*_1_ ≪ 2*N*_0_, *t*_2_ ≫ 2*N*_0_ or *t*_2_ *> t*_1_ ≫ 2*N*_0_ so that the extinction probability Π_0_(*t*_2_ − *t*_1_; *p*) is not negligible. Then due to Eq. (8) and Eq. (10), the integrand

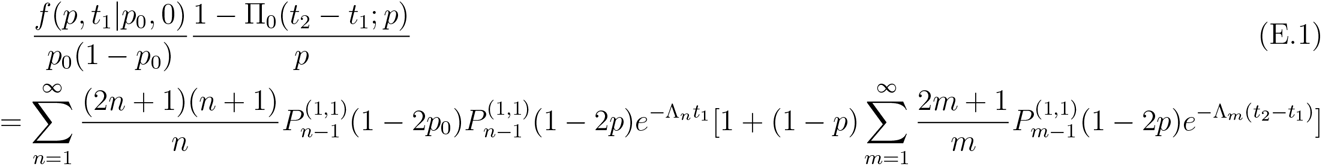

On integrating both sides of the above equation over *p*, we get

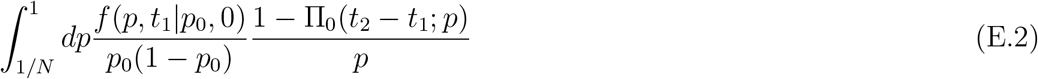

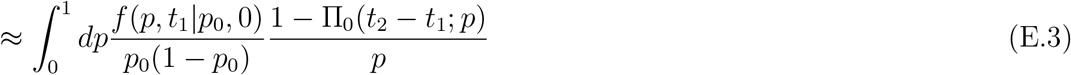

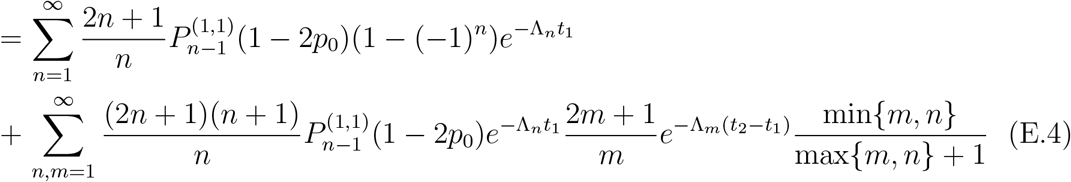

where we have used that

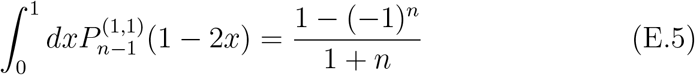

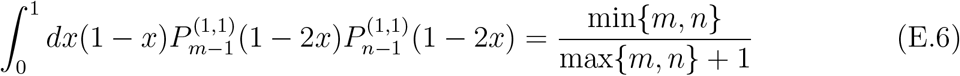

for *m, n* = 1, 2, …. Furthermore, due to Eq. (8) and Eq. (9), the first sum on the RHS of Eq. (E.4) is equal to (1 −Π_0_(*t*_2_; *p*_0_) −Π_1_(*t*_1_; *p*_0_))*/*(*p*_0_(1 − *p*_0_)) on using which we finally arrive at Eq. (36) in the main text.

## Appendix F: Two time correlation function at intermediate times

Here, we consider the time regime *t*_1_ ≪ 2*N*_0_ and 2*N*_0_ ≪ *t*_2_ − *t*_1_ ≪ 2*N*, and develop approximations for the sum on the RHS of Eq. (36) which can be written as *S*_1_ + *S*_2_ where

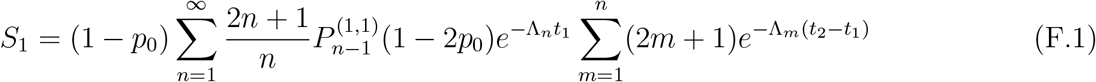

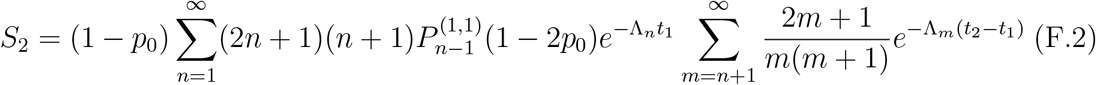

For *t*_2_ − *t*_1_ ≪ 2*N*, as *m* ≫ 1 contributes to *S*_1_ and *S*_2_, we approximate the sum over *m* by an integral as the exact sums do not seem to be doable so that

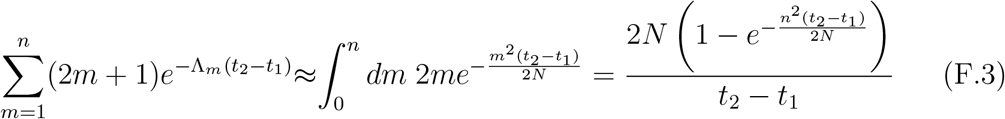

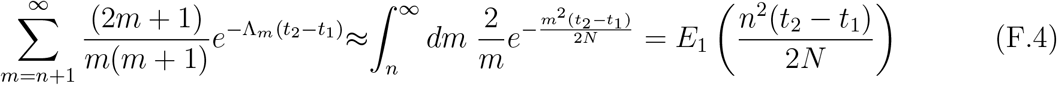

Using the above approximations and Eq. (B.5), for *p*_0_ → 0, we then obtain

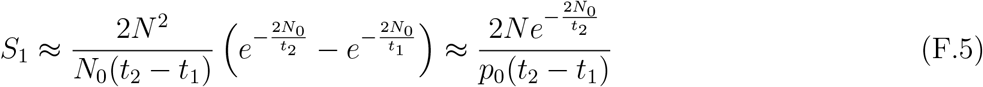

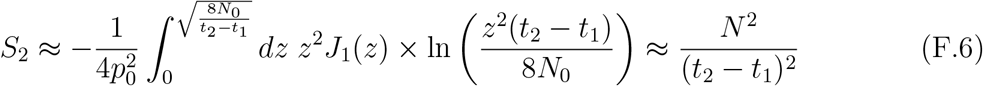

where, for *S*_2_, we have used (6.2.4) of [54] for the power series expansion of the exponential integral. Then, using Eq. (21) and the above approximations in Eq. (36) we finally obtain

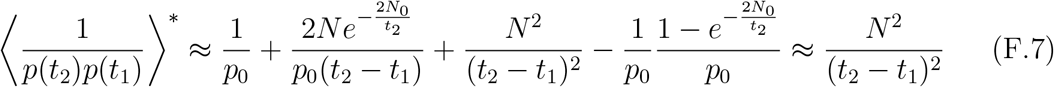

## Appendix G: Exponential decay of the two time correlation function

When *t*_2_ − *t*_1_ ≫ 2*N*, it is sufficient to consider the lowest eigenvalue Λ_*m*_ in the sum over *m* in Eq. (36), but we need to consider the following three cases for the sum over *n*:

i. For *t*_1_ ≪ 2*N*_0_, *t*_2_ − *t*_1_ ≫ 2*N*, from Eq. (36), we can write

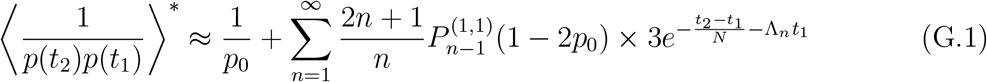

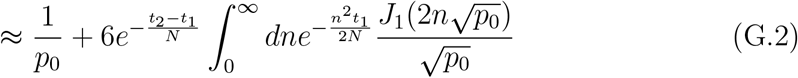

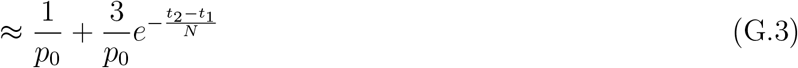

while 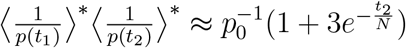 due to Eq. (21) so that

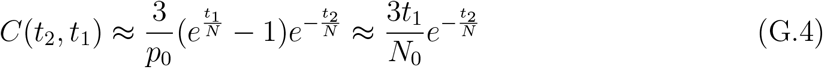
ii. For 2*N*_0_ ≪ *t*_1_ ≪ 2*N, t*_2_ − *t*_1_ ≫ 2*N*, on proceeding as above, we obtain

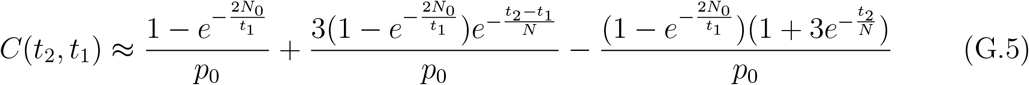

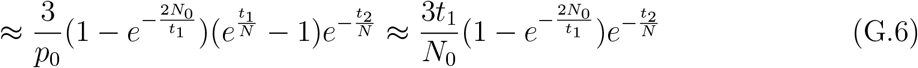
iii. For *t*_1_ ≫ 2*N, t*_2_ − *t*_1_ ≫ 2*N*, it is sufficient to keep terms corresponding to *n* = *m* = 1 in the sum on the RHS of Eq. (36), and we obtain

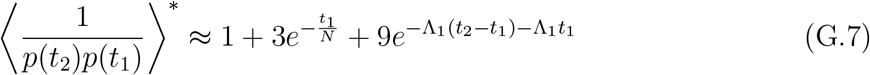

and 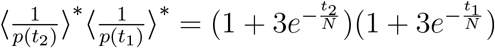 so that

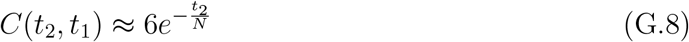

## Appendix H: Distribution of population size at the time of fixation

Here, we discuss the distribution of population size at the time allele 1 fixes in the uncorrelated model. As 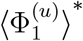 given by Eq. (22) is the cumulative fixation probability by time *t*, on taking its derivative with respect to time, we obtain

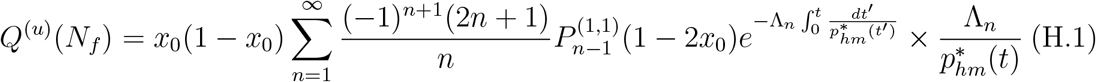

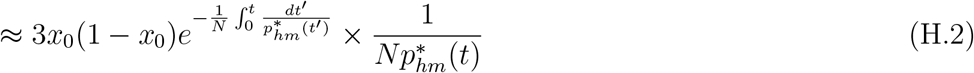

where, due to the discussion in Appendix B for the fixation probability, we have retained only the term corresponding to the lowest eigenvalue. To change the variables from time to population size (which is a random variable), we approximate the relationship through the harmonic mean, 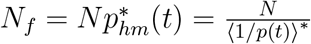 where 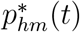 is given by Eq. (17). We then have

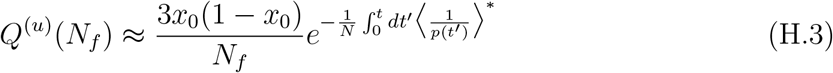

For *t* ≪ 2*N*_0_, from Eq. (21a), we find that *N*_*f*_ ≈ *N*_0_ so that

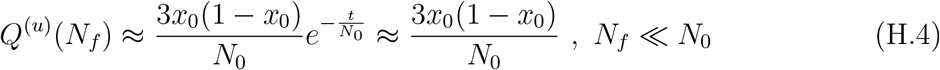

whereas, for 2*N*_0_ ≪ *t* ≪ 2*N*, from Eq. (21b), we can write 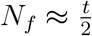 so that

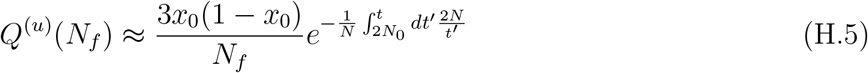

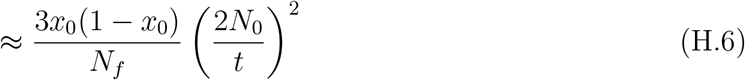

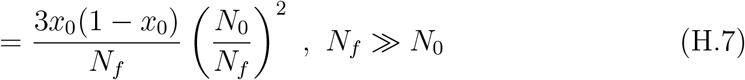

